# Physical activity reduces colorectal cancer risk independent of BMI—A two-sample Mendelian randomisation study

**DOI:** 10.1101/798470

**Authors:** Xiaomeng Zhang, Evropi Theodoratou, Xue Li, Susan M Farrington, Philip J Law, Peter Broderick, Marion Walker, Jessica MB Rees, Richard S Houlston, Ian PM Tomlinson, Harry Campbell, Malcolm G Dunlop, Maria Timofeeva

**Affiliations:** Centre for Global Health Research, Usher Institute of Population Health Sciences and Informatics, The University of Edinburgh, United Kingdom; Colorectal Cancer Genetics Group, CRUK Edinburgh Centre, Institute of Genetics and Molecular Medicine, Western General Hospital, University of Edinburgh, United Kingdom; Division of Genetics and Epidemiology, The Institute of Cancer Research, London, United Kingdom; Colon Cancer Genetics Group, Medical Research Council Human Genetics Unit, Institute of Genetics and Molecular Medicine, Western General Hospital, University of Edinburgh, United Kingdom; Edinburgh Clinical Trials Unit, Centre for Global Health Research, Usher Institute of Population Health Sciences and Informatics, The University of Edinburgh, United Kingdom; Cancer Genetics and Evolution Laboratory, Institute of Cancer and Genomic Sciences, University of Birmingham, Vincent Drive, Edgbaston, Birmingham, United Kingdom

**Keywords:** Physical activity, Body mass index, Colorectal cancer, Mendelian randomisation

## Abstract

**Background:** Evidence from observational studies suggests a protective role for physical activity (PA) against colorectal cancer (CRC) risk. However, it has yet to be established a causal relationship. We conducted a two-sample Mendelian randomisation (MR) study to examine causality between physical activity and CRC risk.

**Methods:** We used common genetic variants associated with self-reported and accelerometer-based physical activity as instrumental variables (IVs) in this MR study. The IVs were derived from the largest available genome-wide association study (GWAS) of physical activity, namely UK Biobank. We analysed the effect of the IVs for physical activity in a large CRC GWAS that included 31 197 cases and 61 770 controls. We applied inverse variance weighted (IVW) method as the main analysis method.

**Results:** Our results demonstrate a protective effect between accelerometer-based physical activity and CRC risk (the outlier-adjusted OR_IVW_ was 0.92 per one standard deviation (SD) increase of accelerometer-base physical activity [95% CI: 0.87-0.98, P: 0.01]). The effect between self-reported physical activity and CRC risk was not statistically significant but was also supportive of an inverse association (the outlier-adjusted OR_IVW_ was 0.61 per 1 SD increase of moderate-to-vigorous physical activity [95%CI: 0.36-1.06, P: 0.08]).

**Conclusions:** The findings of this large MR study show for the first time that objectively measured physical activity is causally implicated in reducing CRC risk. The limitations of the study are that it is based on only two genetic instruments and that it has limited power, despite the study size. Nonetheless, at a population level, these findings provide strong reinforcing evidence to support public health policy measures that encourage exercise, even in obese individuals.

## Introduction

Colorectal cancer (CRC) is the third most common cancer, and the second leading cause of cancer death in the world and the incidence of CRC is projected to increase by 60% by 2040 [1]. More than 1.8 million new cases and 881 000 deaths were estimated to have occurred in 2018 [1]. As a lifestyle risk factor for CRC, convincing evidence from epidemiological studies suggests that physical activity (PA) decreases the risk of colon cancer [2,3]. An umbrella review of 22 anatomical cancer sites including 770 000 cancer cases concluded that there is strong evidence for a protective association between recreational PA and colon cancer [4]. Furthermore, a recent meta-analysis including 17 cohort and 21 case-control studies, found that four domains of PA (occupational activity, recreational activity, transport-related PA and reduced occupational sedentary behaviour) were associated with a lower colon cancer risk, the estimated relative risk (odds ratio) of recreational PA on colon cancer risk was 0.80 (95%CI: 0.71-0.89) and on rectum cancer risk was 0.87 (95%CI: 0.75-1.01) [5].

However, the evidence for the inverse association between PA and CRC is not always consistent. For example, a large prospective study of Norwegian women (n=79 184) did not show an inverse association between PA and CRC risk [6]. Furthermore, causality is not confirmed, mainly due to lack of evidence from randomised clinical trials (RCTs). The observed association could be due to confounding factors (e.g. body weight) or reverse causality. MR can overcome the aforementioned issues, by exploring the effect of the exposure (i.e. PA) on outcome (i.e. CRC) through a genetic instrumental variable (IV) [7]. In a typical MR study, genetic variants are used as an IV, assuming that genetic variants are randomly allocated during gamete formation, similar to a random allocation of RCT participants in intervention or control groups. For the genetic variants to be a valid IVs, three requirements need to be fulfilled: (i) the genetic variants need to be associated with the exposure of interest; (ii) the genetic variants need to be independent of confounders of the exposure-outcome association; and (iii) the genetic variants need to be associated with the outcome only through the exposure. Common variants identified through GWAS can be used as IVs. In this study, we have designed a two-sample summary statistics MR study to explore the causality of the observed association between PA and CRC. As IVs of two different types of PA (moderate-to-vigorous physical activity [MVPA] and mean acceleration vector magnitude [AA]), we used genetic variants that were identified through a large GWAS using UK Biobank data [8].

## Methods

### Data sources

We conducted this MR study by using summary-level data from a large meta-analysis of 15 primary CRC GWAS [9]. Briefly, this published meta-analysis included the *NSCCG-OncoArray GWAS*, the *SCOT GWAS*, The *SOCCS/GS*, *SOCCS/LBC GWAS*, the *UK Biobank GWAS* as well as 10 previously published GWAS studies: UK1, Scotland1, VQ58, CCFR1, CCFR2, COIN, Finnish GWAS, CORSA, DACHS and Croatia [9]. Standard quality-control measures were applied to each GWAS and finally 31 197 cases and 61 770 controls were included in our analysis. All the studies were approved by their respective ethics review committee in accordance with the Declaration of Helsinki.

### Genetic instrument

We conducted MR analyses for two types of continuous PA using common genetic variants (minor allele frequency≥5%) (Additional file 1: Supplementary Table S1) that were derived from a published GWAS study on habitual PA using over 377 000 UK Biobank participants [8]. The two types of PA were: moderate-to-vigorous physical activity (MVPA), which was collected through a self-reported questionnaire for n=377 234 individuals in 2012 and mean acceleration vector magnitude (AA), which was collected from 7 days accelerometer wearing for 91 084 individuals in 2015 [10]. Details of MVPA and AA are described in Additional file 1. This GWAS detected nine Single Nucleotide Polymorphism (SNPs) for MVPA and two SNPs for AA at P<5 ×10 ^−8^. The SNP-based heritability was 5% for MVPA and 14% for AA [8]. If there was linkage disequilibrium, a correlation matrix was added in the statistical model. In addition, we checked the GWAS Catalog [11,12] (https://www.ebi.ac.uk/gwas/home, accessed on 12 July 2019) and other publicly available GWAS consortium data to explore whether the PA instrumental variables were associated with other suspected pleiotropic traits (i.e. body mass index [BMI]).

### Summary data two-sample Mendelian randomisation

Effect estimates of the 11 SNPs on PA were obtained from PA GWAS study and effects of these SNPs on CRC risk were obtained from a meta-analysis of 15 CRC GWASs [9]. The causal effects and the corresponding standard errors of PA on CRC were calculated by using inverse variance-weighted (IVW) method [13]. To conduct a reliable MR analysis, we applied a variety of sensitivity analyses testing different MR assumptions [14]. Specifically, we performed MR– Pleiotropy Residual Sum and Outlier (MR-PRESSO) [15], MR-Robust [16], MR-Egger [17], leave-one-out method [18], mode-based estimate[19] and the median-based method [20]. For the IVW MR [13], we applied fixed effect meta-analysis. We evaluated the heterogeneity between causal effects of each variant (Cochran’s Q statistic) and a P value lower than 0.10 [21] was regarded as statistically significant heterogeneity. MR-PRESSO was applied to identify horizontal pleiotropic outliers [15], MR-robust applied MM-estimation with Tukey’s bisquare function instead of the traditional least squares method, which efficiently limits the contribution of outliers [16], MR-Egger was applied to explore the potential bias introduced by pleiotropy. In particular, when the intercept of MR-Egger differs from zero (at p<0.05), then either directional pleiotropy is indicated or the pleiotropic effects of instruments are correlated with the direct effect (InSIDE assumption) [17]. Finally, we also applied the mode-based estimate, which is consistent when most estimates of identical individual-instrument causal effect derived from valid IVs and weighted median-based method which allows to 50% invalid weights [19,20].

For all MR analyses the P value thresholds were set at 0.05. The statistical analysis of MR was performed on R v3.5.1 with packages ‘MendelianRandomization’ and ‘TwoSampleMR’ [18,22].

### Power calculation

The non-centrality parameter based approach was applied to estimate the power of this study [23]. The self-reported PA related variants explained approximately 0.03% of PA. We fixed the type I error at α<0.05 and applied an assumed true effect estimate of OR=0.80 per 1 SD increase of the PA time. This assumed true effect estimate was obtained from the World Cancer Research Fund meta-analysis of cohort and case-control studies of PA on CRC risk [5]. Power was estimated to be less than 0.10 and F statistic was 14.22 for a sample size of 31 197 CRC cases and 61 770 controls.

## Results

We explored whether horizontal pleiotropy could have biased our results before applying the instrumental variables to perform a two-sample MR. In particular, epidemiological evidence supports an association between increased physical activity and decreased BMI [24] and BMI is associated with CRC risk [25]. Therefore, we checked the GWAS Catalog whether the PA instrumental variables were associated with BMI and examined associations between the PA instrumental variables and BMI with GWAS data from the GIANT consortium [26]. We did not find evidence of association with BMI. Furthermore, a meta-analysis on subgroup analysis of the PA-CRC risk relation indicated that PA is associated with CRC risk in both high and low BMI groups [27]. We also checked the GWAS catalog to identify if any of the PA genetic instrumental variables was associated with other traits. We found that rs55657917 and rs1043595 were associated with neuroticism and intelligence respectively [28,29].

Furthermore, a recent MR study supported a protective relationship between PA and major depressive disorder [30]. However, no epidemiological evidence links depression with an increased CRC risk [31].

### Self-reported MVPA and CRC

The associations of the nine MVPA SNPs with MVPA and with CRC are presented in Additional file 1: Supplementary Table S2. Based on the MR results, no statistically significant causal effect was detected between self-reported MVPA and CRC risk (Table 1). However, the result for the IV of MVPA SNPs was suggestive of an association after removing the outlier detected by MR-PRESSO and leave-one-out method (rs429358). The outlier-adjusted OR of IVW MR was 0.61 for CRC risk per 1 SD increase of MVPA (95%CI: 0.36-1.06, P=0.08) (Table 1, Figure 1). Result from the MR-robust analysis also suggested a potential PA-CRC association (95%CI: 0.37-1.01, P=0.054) while other sensitivity analyses like the median-based method and mode-base estimate did not report significant causal effect (Table 1). The intercept of MR-Egger regression did not identify any horizontal pleiotropy and/or violation of the InSIDE assumption (P=0.51) (Table 1). Similarly, after removing the outlier, Q statistic did not indicate nominal heterogeneity (P=0.397 vs P=0.002). The outlier we detected mapped on the APOE gene, which has been found to be associated with a number of other phenotypic traits [32,33]. In addition, through our search in the GWAS Catalog, we identified rs1043595 to be related to cognitive function and intelligence [29]. However, removing this SNP did not change the result.

**Table 1.** Results of Two-sample Mendelian randomisation analysis for MVPA and AA

|  |  |  |  |  |  |  |  |  |  |  |  |  |  |
| --- | --- | --- | --- | --- | --- | --- | --- | --- | --- | --- | --- | --- | --- |
|  | MVPA |  |  |  |  | MVPA<sup>C</sup> |  |  |  |  | AA |  |  |
|  | Causal effects<br>(95% CI) | P | P<sub>int</sub><sup>a</sup> | P<sub>het</sub><sup>b</sup> |  | Causal effects<br>(95% CI) | P | P<sub>int</sub><sup>a</sup> | P<sub>het</sub><sup>b</sup> |  | Causal effects<br>(95% CI) | P | P<sub>het</sub><sup>b</sup> |
| Main analysis |  |  |  |  |  |  |  |  |  |  |  |  |  |
| IVW | 0.92 (0.38,2.21) | 0.85 | / | 0.002 |  | 0.61 (0.36,1.06) | 0.08 | / | 0.40 |  | 0.92 (0.87,0.98) | 0.01 | 0.16 |
| Sensitivity analyses |  |  |  |  |  |  |  |  |  |  |  |  |  |
| MBE | 0.53 (0.20,1.38) | 0.19 | / | / |  | 0.48 (0.16,1.41) | 0.18 | / | / |  | 0.92 (0.86,0.98) | 0.01 | / |
| Weighted<br>median | 0.67 (0.32,1.39) | 0.28 | / | / |  | 0.59 (0.29,1.21) | 0.15 | / | / |  | / | / | / |
| MR-Egger | 1.60 (0.05,56.24) | 0.80 | 0.75 | 0.001 |  | 0.30 (0.03,2.64) | 0.28 | 0.50 | 0.34 |  | / | / | / |
| MR-Robust | 0.72 (0.30,1.73) | 0.46 | / | / |  | 0.61 (0.37,1.01) | 0.05 | / | / |  | 0.92 (0.89,0.95) | 3.97E-07 | / |
| MR-PRESSO | 0.92 (0.38,2.21) | 0.85 | / | / |  | 0.60 (0.35,1.05) | 0.11 | / | / |  | / | / | / |
| MVPA: Moderate-to-vigorous physical activity, <sup>C</sup>MVPA: removed rs429358 base on the result of MR-PRESSO and leave-one-out methods, AA: Average acceleration, CI: Confidence interval, <sup>a</sup>P<sub>int</sub>: P value for the intercept of MR-Egger's test, <sup>b</sup>P<sub>het</sub>: P values of $\chi^2$ Q test for heterogeneity, IVW: Inverse variance-weighted, MBE: Mode-based estimate, MR-PRESSO: MR-Pleiotropy Residual Sum and Outlier | | | | | | | | | | | | | |
| AA only has two available genetic instruments, we applied limited number of methods. |  |  |  |  |  |  |  |  |  |  |  |  |  |

**Figure 1.**
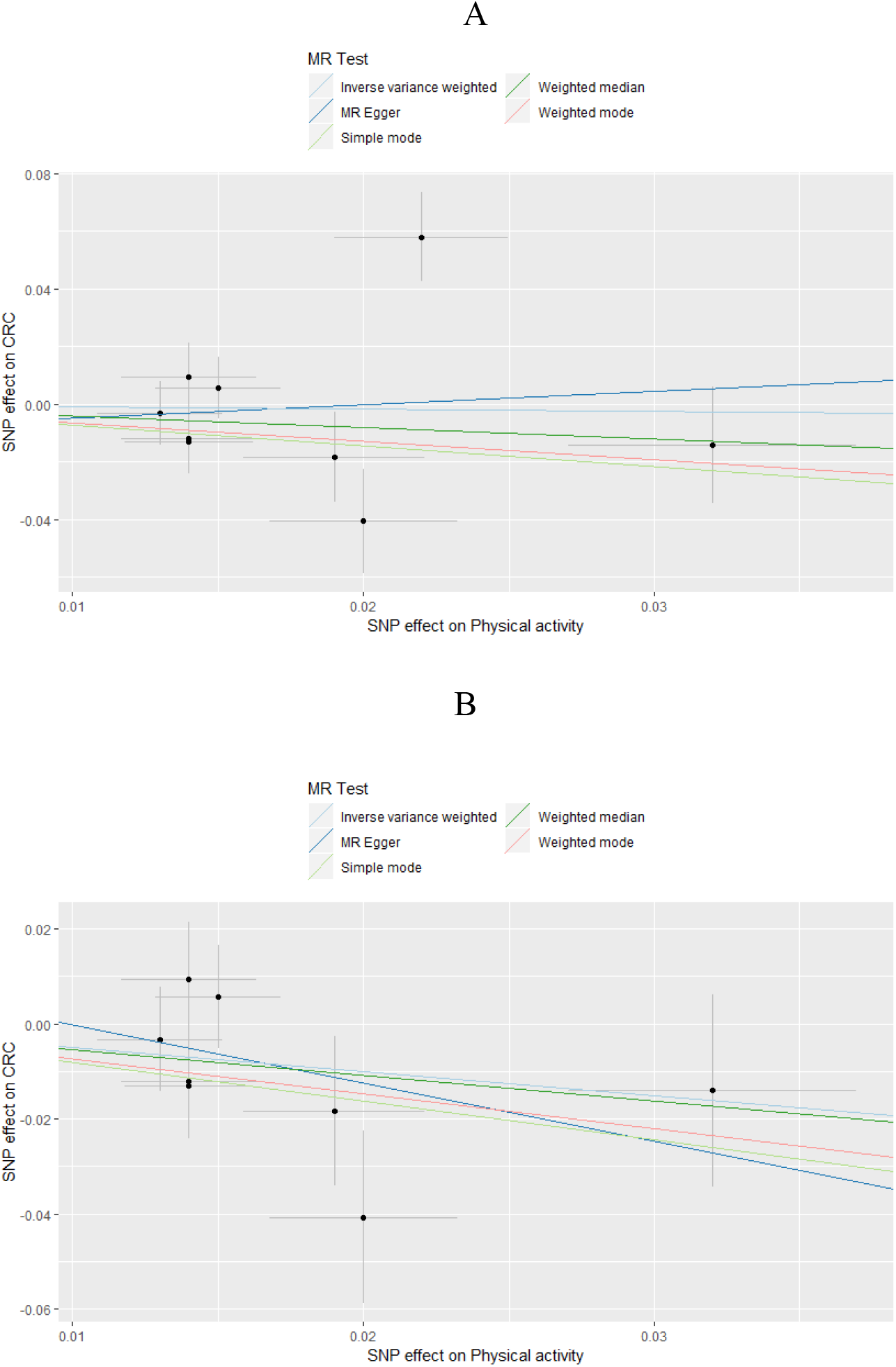
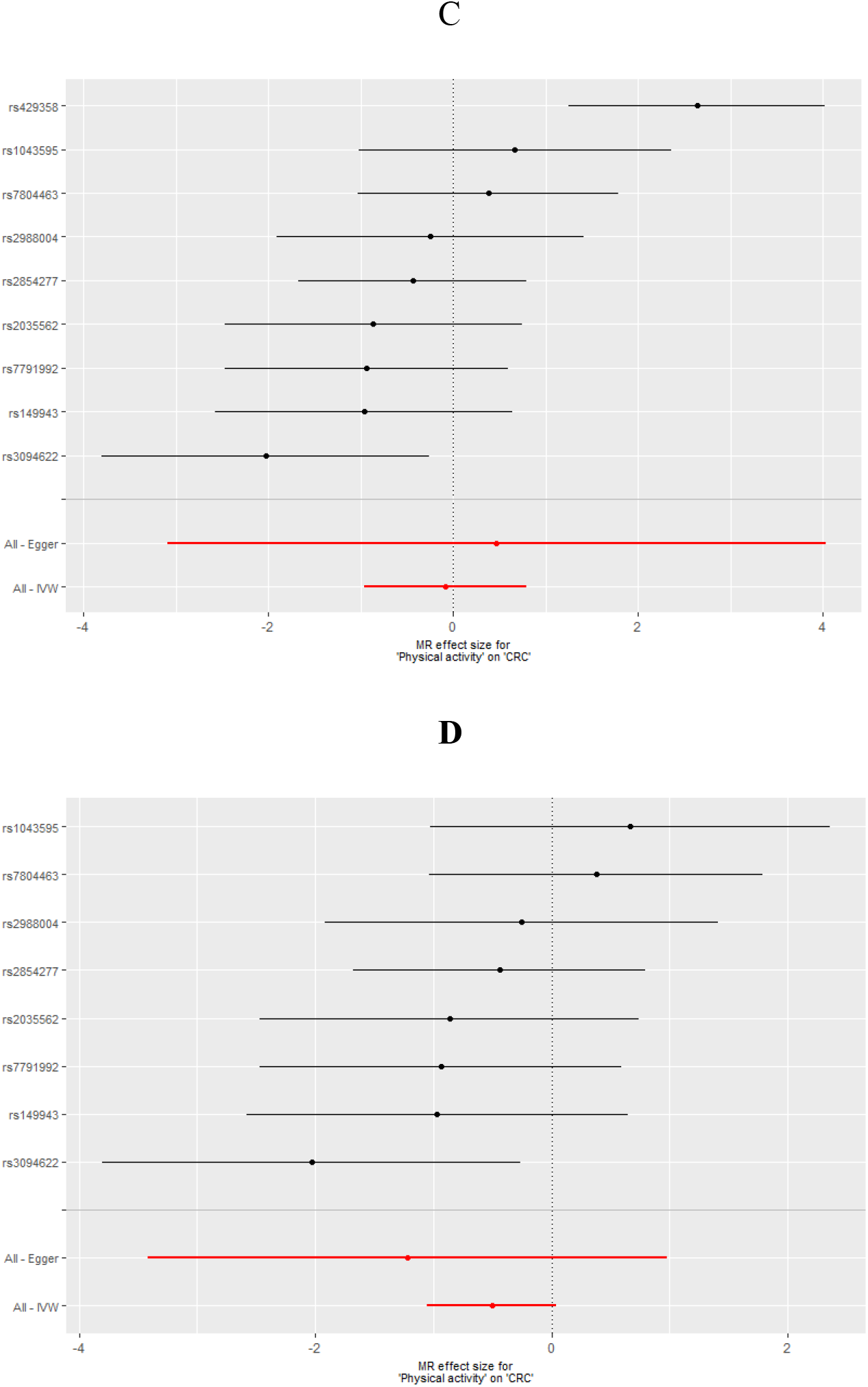
Visualisation of MR analysis of the effect of self-reported physical activity on colorectal cancer risk. A: Scatter plot without removing outlier SNP detected by MR-PRESSO/ leave-one-out method B: Scatter plot after removing outlier SNP rs429358 detected by MR-PRESSO/ leave-one-out method C: Forest plot without removing outlier SNP detected by MR-PRESSO/ leave-one-out method D: Forest plot after removing outlier SNP rs429358 detected by MR-PRESSO/ leave-one-out method

### Accelerometer-based physical activity and CRC

The effect estimates of each genetic variant on AA and on CRC is listed in Additional file 1: Supplementary Table S2. Evidence for causal association was detected between accelerometer based physical activity and CRC risk by using the two SNPs as genetic instruments (Table 1, Figure 2). In particular, the OR of IVW MR was 0.92 (95% CI: 0.87-0.98, P=0.01) (Table 1).

**Figure 2.**
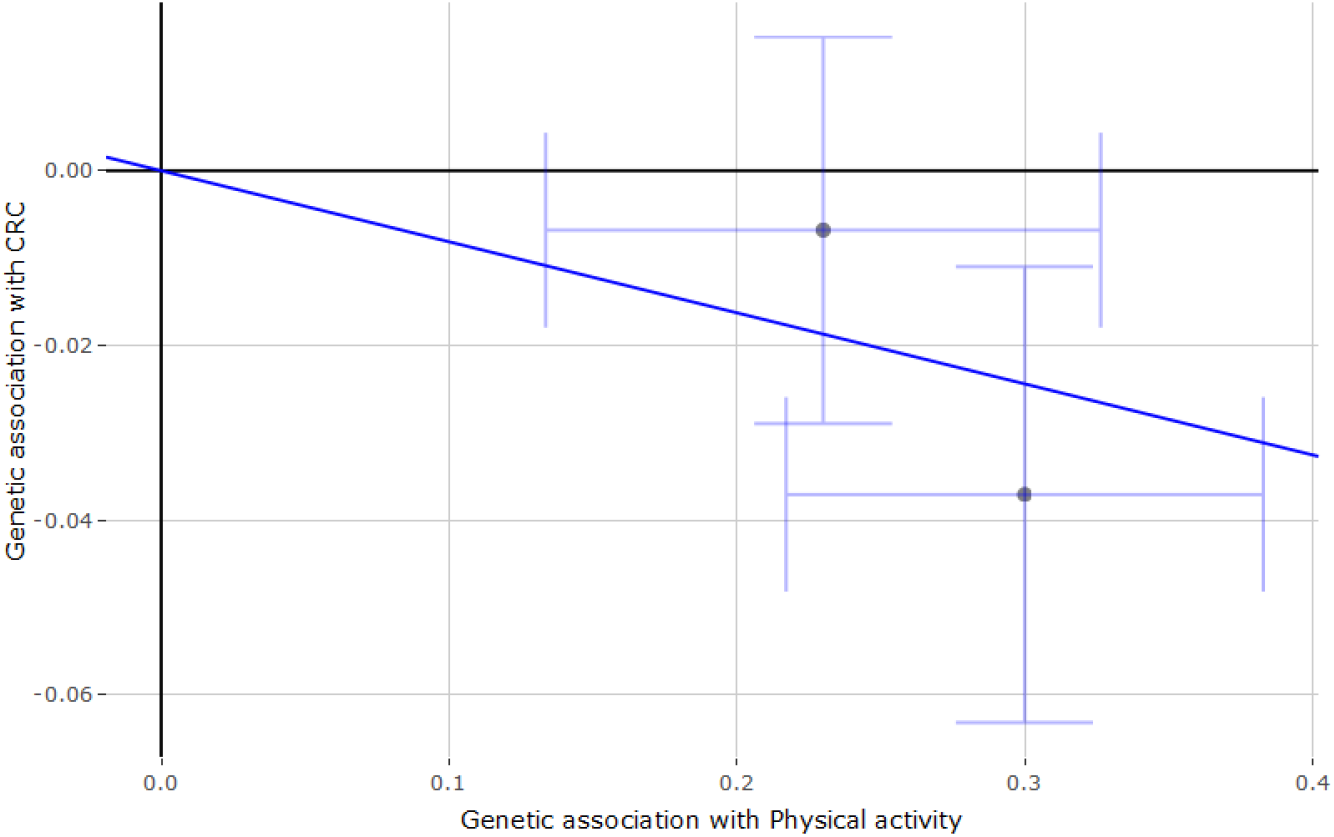
Scatterplot of SNP effects on accelerometer-based physical activity versus the effects on colorectal cancer. Scatterplot and regression line conducted by using IVW method with effects estimate and its 95% confidence interval on SNPs-PA and SNPs-CRC associations

As sensitivity analyses, both mode-based estimate and MR-robust supported the causal association between accelerometer-based PA and CRC risk (Table 1). Due to the small number of genetic variants used for IV, we could not apply other sensitivity analyses. The Q statistic suggested no heterogeneity (P=0.16).

## Discussion

CRC is a common cancer with significant morbidity and mortality [1] and with an increasingly incidence rate in younger ages [34]. Prospective cohort, case-control and cross-sectional observational studies support an inverse association between PA and CRC risk [3,35,36]. The World Cancer Research Fund Network also reported a convincing evidence on the inverse association between PA and colon cancer from prospective observational studies [37]. However, there is a lack of RCTs that could further investigate the possible causal role of low levels of PA in CRC development. In this study, genetic variants from a very large GWAS were employed as instruments to explore the causal effect between PA and CRC. By applying two-sample MR, we observed a causal effect of accelerometer-based PA on CRC and a suggestive association between self-reported PA and CRC [14]. The MR results for both types of PA were similar to the results from the meta-analyses of observational studies [5].

The causal effect estimate between accelerometer-based PA and CRC suggests a slight decrease of CRC risk per 1 SD accelerometer-based PA increase. Although there is no standard method to transform milli-gravities to energy expenditure, 1 SD change of accelerometer-based PA (8.14 milli-gravities or 0.08m/s^2^) approximates to about 3 metabolic equivalent task (MET)-hour/day [10]. This means every day, if people replace sedentary behaviour with 26 to 45 minutes of moderate PA (such as 45 minutes 10 mph bicycling or 26 minutes hiking) or with 13 to 26 minutes of vigorous PA (such as 23 minutes of swimming or 13 minutes of running), their risk of CRC will decrease by 8% [38].

Furthermore, causal effect estimate of MVPA against CRC risk indicated a potential association between self-reported PA and CRC risk, which suggests that with 1 SD increase of moderate-to-vigorous PA CRC risk decreases by 39%. One SD of MVPA is around 4.96 MET-hour/day. This result suggests that every day, if people replace sedentary behaviour with 43 to 74 minutes moderate PA (such as 74 minutes 10 mph bicycling or 43 minutes hiking) or with 22 to 43 minutes of vigorous PA (such as 37 minutes of swimming or 22 minutes of running), their risk to develop CRC may have a moderate decrease by 39% [38]. As we know, self-reported PA tends to be overestimated and that may result in an overestimation of the effect on CRC risk.

Diverse biologic mechanisms have been proposed to support the observed inverse association between PA and CRC such as the beneficial effect of PA on bowel transit time [39,40], the immune system reactions [41], the metabolism of bile acid, the better insulin sensitivity[42] and the reduction of prostaglandin E2 levels in colonic mucosa [43]. Evidence from RCTs supports that PA can reduce the bowel transit time and therefore reduce the time of contact between carcinogens and colonic mucosa [39,40]. The increase of prostaglandin synthesis can also promote the intestinal peristalsis and hence reduce transit time [44]. In addition, prostaglandin E2 can promote tumour generation directly or through its multifaceted effects on inflammation [45]. Physical activity also relates to a lower concentration of bile acid, which is an essential mediator of the cholesterol mechanism and the lower bile acid concentration is associated to a lower blood triglycerides [46]. Through this pathway, the effect between PA and CRC risk could be through obesity related markers. In addition, regular moderate PA may have a benefit on natural cytotoxicity and T-lymphocyte proliferation, on reducing the production of pro-inflammatory cytokines and on increasing the count of T-cells, B-cells and Immunoglobulins [41].

### Strengths and limitations

One of the strengths of this study was that we explored both subjective and objective measures of PA (self-reported PA and accelerometer-based PA). Previous studies showed that there are discrepancies between self-reported PA and accelerometer-based PA [47,48]. Compared to self-reported PA which is easy to be affected by body health and emotional health, the accelerometer-based PA explains 44-47% variance of energy expenditure [49]. On the other hand, self-reported PA tends to overestimate the time of doing PA in the general population [50]. Furthermore, the application of instrumental variable methods provides a new insight on PA research. The genetic instruments for PA were detected from a UK-biobank GWAS which is a large cohort among people aged 40 to 70 years old.

This study has also several limitations. First, although we cited the largest available GWAS for PA, the detected SNPs for self-reported PA only explain a low proportion of the PA variance (0.03%). Even though the calculated F statistic (F statistic=14.22) reached a previously suggested threshold level for avoiding weak instrument bias [51], the instrumental variables explain only a limited proportion of overall variance in PA. As a result, our analysis was underpowered (<0.8). Second, with a 14% heritability for accelerometer-base PA, the low variance of genetic instruments may imply that the current discovered SNPs cannot be considered as powerful proxies for physical activity. Furthermore, although we applied the most up-to-date MR methods, we cannot completely rule out any potential horizontal pleiotropy until we know all the biological functions for each SNP. Third, in two-sample MR analysis, weak instrument bias is in the direction of null while the partial overlapping data between exposure and outcome from UK-biobank may dis-equilibrate the direction of null [52]. The number of CRC cases from UK-biobank is 6360 which accounts for about 6.8% of the total participants in this analysis. Even if all of the 6360 patients overlapped, this proportion is low and will not affect the effect estimation between PA and CRC risk [52].

Fourth, because we only have two SNPs as IVs for accelerometer-based PA, the other MR methods or sensitivity analyses cannot be applied. Finally, the data we have were not available to analyse the association between PA and colon and rectal cancer separately, even though evidence from a European multinational cohort study showed that PA was associated with proximal colon cancer and distal colon cancer risk but not with rectal cancer risk [53].

## Conclusions

The incidence of CRC is increasing among younger adults, with some experts suggesting that it could be due to higher prevalence of CRC risk factors (including obesity) in the younger populations [34,54]. Results of this study indicate a causal role of objectively measured PA in CRC risk independent of BMI. Therefore, promotion of physical activity could probably result in the decrease of CRC incidence, even in the younger populations or for obese individuals.

## Acknowledgements

We acknowledge the excellent technical support from Stuart Reid. We are grateful to Donna Markie and all those who continue to contribute to recruitment, data collection, and data curation for the Study of Colorectal Cancer in Scotland studies. We acknowledge that these studies would not be possible without the patients and surgeons who take part. We acknowledge the expert support on sample preparation from the Genetics Core of the Edinburgh Welcome Trust Clinical Research Facility.

## Authors’ contributions

MGD, MT and ET conceived this study. XZ, MT, ET, JR and XL designed the methodology. XZ conducted data analysis with the supervision from MT and ET. XZ drafted the manuscript, conducted data interpretation with ET, MT and MGD. XL, SF, PL, PB, MW, JR, RH, IT and HC contributed to the manuscript drafting and revision. XZ and ET contribute equally to this paper.

## Appendix A. Supplementary data

