## Supplemental data for "Physical activity reduces colorectal cancer risk independent of BMI—A two-sample Mendelian randomisation study"

**Description of MVPA and AA**

UK Biobank collected physical activity data during work and leisure time through self-reported questionnaire and 7 days Accelerometer wearing [1, 2]. For self-reported moderate PA (MPA) and vigorous PA (VPA), participants were asked: ‘In a typical WEEK, on how many days did you do 10 min or more of moderate physical activities like carrying light loads, cycling at normal pace? (Do not include walking)’ and ‘In a typical WEEK, how many days did you do 10 min or more of vigorous physical activity? (These are activities that make you sweat or breathe hard such as fast cycling, aerobics, heavy lifting)’ respectively. For each of these questions, those who indicated 1 or more such days were then asked ‘How many minutes did you usually spend doing moderate/vigorous activities on a typical DAY’. Moderate-to-vigorous PA (MVPA) was calculated by taking the sum of total minutes/week of MPA multiplied by four and the total number of VPA minutes/week multiplied by eight. In the UK Biobank, about 103712 datasets collected from participants wearing an Axivity AX3 wrist-worn accelerometer for seven days. We included one measure from accelerometer wearing: the mean acceleration vector magnitude (AA). The unit for physical activity vector magnitude is milli-gravities (mg) and the calculation described in a UK Biobank study [2]. The details of MVPA and AA were summarized in (Supplementary Table S1).

**Supplementary tables**

Supplementary Table S1 Genetic variants associated with two continuous physical activity types recorded from the UK Biobank study.

| Category | Summary | rsid | Chr | Closest gene | Position | EA | EAF |
|---|---|---|---|---|---|---|---|
| UK Biobank: The mean acceleration vector magnitude **(AA)** (milli-gravities) | Mean=27.98; Median=27.03; SD=8.14; n=91,084 | rs55657917 | 17 | CRHR1 | 43,844,560 | T | 0.78 |
| | | rs59499656 | 18 | RIT2/SYT4 | 40,768,309 | A | 0.66 |
| UK Biobank: Moderate-to-vigorous physical activity **(MVPA)** (MET-minutes/week) | Mean=1650; Median=960; SD=2084; n=90,667 | rs429358 | 19 | APOE | 45,411,941 | T | 0.85 |
| | | rs2035562 | 3 | CADM2 | 85,056,521 | A | 0.33 |
| | | rs7791992 | 7 | C7orf72/SPATA48 | 50,237,784 | C | 0.41 |
| | | rs2988004 | 9 | PAX5 | 37,044,388 | T | 0.56 |
| | | rs1043595 | 7 | CALU | 128,410,012 | G | 0.72 |
| | | rs7804463 | 7 | EXOC4 | 133,447,651 | T | 0.53 |
| | | rs149943 | 6 | ZNF165 | 28,002,388 | G | 0.85 |
| | | rs3094622 | 6 | RPP21 | 30,327,952 | A | 0.86 |
| | | rs2854277 | 6 | HLA-DQB1 | 32,628,084 | C | 0.92 |

Genetic variants for PA were extracted from a physical activity GWAS study [1].
MPA: Moderate physical activity, VPA: Vigorous physical activity, SD: Standard deviation, EA: Effect allele, EAF: Effect allele frequency

| Supplementary Table S2 Summary of associations between genetic variants and physical activity/colorectal cancer risk | | | | | | | | | | | |
|---|---|---|---|---|---|---|---|---|---|---|---|
| **Category** | **Chr** | **Position** | **rsid** | **Closest gene** | **EA for PA/CRC** | **EAF** | **Beta.PA** | **SE.PA** | **P.PA** | **Beta.CRC** | **SE.CRC** |
| AA | 17 | 43844560 | rs55657917 | CRHR1 | G (T) | 0.78 | 0.300 | 0.042 | 5.0E-12 | -0.037 | 0.013 |
| AA | 18 | 40768309 | rs59499656 | RIT2/SYT4 | T (A) | 0.66 | 0.230 | 0.038 | 2.4E-09 | -0.007 | 0.011 |
| MVPA | 19 | 45411941 | rs429358 | APOE | C (T) | 0.85 | 0.022 | 0.003 | 6.1E-13 | 0.058 | 0.016 |
| MVPA | 3 | 85056521 | rs2035562 | CADM2 | G (A) | 0.33 | 0.014 | 0.002 | 3.9E-09 | -0.012 | 0.011 |
| MVPA | 6 | 28002388 | rs149943 | ZNF165 | A (G) | 0.85 | -0.019 | 0.003 | 2.2E-09 | 0.018 | 0.016 |
| MVPA | 6 | 30327952 | rs3094622 | RPP21 | G (A) | 0.86 | -0.020 | 0.003 | 1.4E-09 | 0.041 | 0.018 |
| MVPA | 6 | 32628084 | rs2854277 | HLA-DQB1 | T (C) | 0.92 | -0.032 | 0.005 | 2.6E-10 | 0.014 | 0.020 |
| MVPA | 7 | 128410012 | rs1043595 | CALU | A (G) | 0.72 | -0.014 | 0.002 | 4.3E-09 | -0.009 | 0.012 |
| MVPA | 7 | 133447651 | rs7804463 | EXOC4 | C (T) | 0.53 | -0.015 | 0.002 | 1.2E-11 | -0.006 | 0.011 |
| MVPA | 7 | 50237784 | rs7791992 | C7orf72/SPATA48 | A (C) | 0.41 | 0.014 | 0.002 | 5.7E-10 | -0.013 | 0.011 |
| MVPA | 9 | 37044388 | rs2988004 | PAX5 | G (T) | 0.56 | 0.013 | 0.002 | 4.1E-09 | -0.003 | 0.011 |
| PA: Physical activity, MVPA: Moderate-to-vigorous physical activity, AA: Average acceleration, EA: Effect allele, EAF: Effect allele frequency, CRC: Colorectal cancer, SE: Standard error<br>Associations between genetic variants and PA were extracted from a physical activity GWAS study [1], Associations between PA genetic variants and CRC risk were extracted from a meta-analysis of 15 primary CRC GWAS [3]. | | | | | | | | | | | |
